# Ancestral diet leads to dynamic transgenerational plasticity for five generations in *Drosophila melanogaster*

**DOI:** 10.1101/273144

**Authors:** Carmen Emborski, Alexander S Mikheyev

## Abstract

Ancestral exposures can influence phenotypic expression in subsequent generations, which influence diverse biological processes ranging from phenotypic plasticity to obesity. Currently, most transgenerational studies work under the assumption of transgenerational response stability and reproducibility through time and across exposure differences, relying on short-term (i.e. 2-3 generations) single-exposure experiments. Yet, little evidence exists in the literature to validate this assumption, leaving the consistency and reliability of interpretations in question. Additionally, as most studies have focused on proximal mechanistic (‘how’) rather than ultimate evolutionary (‘why’) questions, the interpretations of observed responses and broader evolutionary implications remain unclear. In the current study, we begin to address these gaps by analyzing the transgenerational effects of three dietary sugar concentrations (*i.e.* no, low, and high) relative to controls on body composition (*i.e.* whole body fat and sugar concentrations) and reproduction (*i.e.* lifetime fitness) over five generations in both males and females. We found that the changes in ancestral diet led to complex transgenerational body composition and fitness fluctuations relative to control offspring responses, despite the conformity of the treatments to a control diet in the F1-F5 generations. Interestingly, the direction of response frequently changed from generation to generation, and as a function of ancestral exposures and sex. These fluctuating response findings have not been documented previously, and were broadly consistent in both our pilot and main experiments. Our results highlight the dynamic and multifaceted nature of transgenerational plasticity, and provide some of the first evidence that transgenerational response stability may not be universally valid.

**Summary Statement:** Alterations in ancestral dietary sugar transgenerationally influences body composition and fitness in Drosophila melanogaster, leading to fluctuating responses relative to controls over five generations following exposure.

## Introduction

Until recently, the dominant view of phenotypic change from generation to generation followed a combination of Mendel’s and Darwin’s views on natural selection through genetic variation (Laland et al., 2015). Increasingly, this view has been challenged by evidence documenting a parent’s ability to transmit phenotypic traits to their offspring, which occur not just at conception but also via subsequent developmental interactions between parent and offspring (Laland et al., 2015, Bonduriansky and Day, 2009). An increasing number of studies have shown that an ancestor’s exposure can influence the condition and response of multiple subsequent generations even in the absence of current exposures (i.e. transgenerational phenotypic plasticity) (Bonduriansky, 2012, Jablonka and Raz, 2009, Uller, 2008), which has been found to occur across a range of organisms (Xu et al., 2014, Tracey et al., 2013, Chamorro-Garcia et al., 2013, Valtonen et al., 2012, Rechavi et al., 2011, Painter et al., 2008). When ancestral experience affects offspring responses at least two or three generations after exposure, where the specific number of generations analyzed depends on the species’ reproductive strategy (namely when germ-line cells differentiate in development) (Buescher et al., 2013, Dew-Budd et al., 2016, Xia and De Belle, 2016, Dunn and Bale, 2011, Tauffenberger and Parker, 2014, Benyshek et al., 2006, Skinner, 2008), these effects are considered transgenerational.

Transgenerational inheritance and plasticity has received considerable interest in the literature (Dunn and Bale, 2011, Ayyanath et al., 2013, Buescher et al., 2013, Pentinat et al., 2010, Cropley et al., 2016). From these studies, results primarily describe one of three findings: stable, delayed, or diminished effects. Stable effects occur when responses are consistently elevated or depressed across multiple subsequent generations (Remy, 2010, Rechavi et al., 2014). Delayedtransgenerational effects occur when phenotypic changes are not present immediately (e.g., skipping a generation before influencing a response)(Benyshek et al., 2006). Diminishing effects occur when responses are seen immediately, remaining steady for one or a few generations after exposure, and then taper back to the level of control by the last (typically second or third) generation analyzed, thus suggesting no true transgenerational effect (Sagi-Schwartz et al., 2008, Armitage et al., 2007). Less commonly, compensatory effects have also been described (Hoile et al., 2011, Shama and Wegner, 2014, Vyssotski, 2011). Here, transgenerational inheritance is also seen, but the signals transferred to and subsequent phenotype(s) of offspring are intergenerationally adjusted based on the interaction between the parental phenotype and interpreted environment.

Although stable transgenerational responses are often the gold standard of what is considered transgenerational effects, they are actually fairly uncommon in the literature (Remy, 2010, Rechavi et al., 2014, Ashe et al., 2012). Conversely, delayed and diminishing responses are quite common for transgenerational studies (J Marshall and Uller, 2007, Gluckman et al., 2005, Nystrand et al., 2016, Xia and De Belle, 2016, Dew-Budd et al., 2016, Buescher et al., 2013, Dunn and Bale, 2011, Pentinat et al., 2010, Franklin et al., 2010, Walsh et al., 2015). Unfortunately, these aforementioned interpretations are frequently based on studies with short-time scales and single-exposure changes, limiting the scope of inference. For example, as transgenerational studies frequently only analyze a single extreme alteration of the stressor (e.g., a single dietary modification treatment of increased sugar or fat concentrations), the consistency of transgenerational responses across differing diets or concentrations of dietary modifications is generally unknown (Kovalchuk, 2012). This is particularly true as few studies analyze multiple concentrations or stressors under the same experimental conditions or within the same study (Matzkin et al., 2013, Wu and Suzuki, 2006, Musselman et al., 2011, Benyshek et al., 2006, Dunn and Bale, 2011).

Additionally, as studies generally only analyze the minimum timeframes needed to document transgenerational inheritance (e.g., 2-3 generations following exposure) (Xia and De Belle, 2016, Shama and Wegner, 2014, Manikkam et al., 2013, Tracey et al., 2013, Chamorro-Garcia et al., 2013, Dunn and Bale, 2011, Buescher et al., 2013, Tauffenberger and Parker, 2014, Franklin et al., 2010, Cropley et al., 2016), the stability of offspring responses over the course of generational time is not well-documented. Taken together, although delayed and diminished transgenerational study interpretations and conclusions appear to work under the assumption of transgenerational response stability and reproducibility through time and across exposure differences, there is little evidence in the literature to support it (Kovalchuk, 2012). Thus, the consistency and reliability of previous study interpretations and conclusions are left in question.

Additionally, a major factor limiting the functional insights that transgenerational studies can provide is the lack of evolutionary insight, particularly given that most studies focus exclusively on the proximal mechanistic ‘how’, rather than the ultimate evolutionary ‘why’. For example, due to the increasing prevalence of non-communicable disease such as metabolic disease (i.e. diabetes and obesity), many inter- and trans-generational studies have focused exclusively on metabolic or body composition effects (Chamorro-Garcia et al., 2013, Buescher et al., 2013, Valtonen et al., 2012, Gluckman et al., 2007, Ost et al., 2014, Painter et al., 2008, Pentinat et al., 2010, Wu and Suzuki, 2006, Xu et al., 2014, Cropley et al., 2016). Though body composition provides a snapshot into an individual’s current state, it is difficult to clearly define whether these alterations significantly affect an organism’s survival or reproductive abilities, which would provide better insight into its adaptive evolutionary context. Lifetime reproductive fitness has a well-documented relationship with body composition (Pasquali, 2003, Ruff et al., 2013, Xu et al., 2011), yet fitness measurements are rarely included in health related studies, giving these studies virtually no adaptive context. Therefore, in addition to analyzing proximate endpoints like body composition, analyzing lifetime reproductive fitness may help give a more measureable proxy for wellness as it relates to disease, such as metabolic disorders, and provide a better evolutionary context of any ancestral effects.

Taken together, the simplicity of study designs and inconsistency between findings, whether due to measuring 1) too few of stressor concentrations or magnitudes, 2) too short time periods, or 3) different response endpoints, has made it difficult to find ultimate patterns of responses and better define the functional and evolutionary value of transgenerational inheritance. In the current study, we attempt to overcome some of these limitations by using a tractable model system, *Drosophila melanogaster* (i.e. fruit flies), to create a more powerful experimental design. We test the hypothesis that a single generation exposure to three different concentrations of a sugar can alter offspring body composition and reproductive fitness for at least five subsequent generations, as compared to controls.

Fruit flies are a flexible and robust model system to study diet-induced inheritance due to their short and well-defined life cycles, simple and easily altered diet, and broad metabolic, digestive, and regulatory similarities to mammals and other eukaryotes (Lemaitre and Miguel-Aliaga, 2013). In particular, fruit flies offer a feasible opportunity to run longer and more complex studies, including direct measurements of lifetime fitness. For example, by analyzing three different levels of ancestral sugar exposure (i.e. no-, low-, or high-sugar treatment diets) relative to a control, we are better able to understand the consistency of transgenerational responses between differing exposures. Additionally, by studying body composition and fitness responses over five generations after exposure, we are able to better document their longer-term stability or instability of both positive (i.e. health) and adaptive (i.e. reproductive success) endpoints.

We found novel response patterns, where responses fluctuated from generation to generation relative to control and differed depending on the treatment concentration and sex. Our findings contradict commonly invoked explanations described in simpler transgenerational studies. Overall, we were able to delve more deeply into the transgenerational influence that ancestral experience has on offspring phenotypes and show more comprehensive transgenerational patterns that may be influencing phenotypic plasticity.

## Results

### Sex-specific on metabolite concentrations

Metabolite concentrations significantly differed between sexes when all treatments and weight were taken into account (supplementary table 1), which was the case for glucose (0.026), trehalose (*p* = <0.0001), glycogen (*p* = 0.00024), and triglycerides (*p* = 0.000792). Additionally, treatment effects in females generally showed a larger frequency of significant differences relative to controls, as compared to males.

### Dietary treatments affect metabolite concentrations and fitness (F0)

As expected, the three dietary sugar exposures generally led to corresponding metabolite concentrations in the exposure (F0) generation (figure 2) (supplementary figures 2 and 3). Specifically, sugar and fat concentrations of F0 NSD flies were significantly lower than CD flies in both female and male flies (*p =* <0.001 for all metabolites). In the LSD treatment, female metabolite concentrations were also significantly lower than controls for sugar and fat concentrations (glucose (*p* = 0.029), glycogen (*p* = <0.001) and triglycerides (*p* = <0.001)), whereas males only differed significantly from controls in fat concentrations (*p=* 0.001). As expected, all LSD metabolite concentrations were higher than NSD metabolite concentrations in both sexes. In the HSD flies, both sexes significantly differed in glycogen (p = 0.02) and triglyceride concentrations (*p* = <0.01), though no significant differences were found in glucose or trehalose concentrations. In all three treatments, F0 fitness measurements did not significantly differ from controls (NSD: *p* = 0.417; LSD: *p* = 0.466, *p* = 0.981).

**Figure 1:**
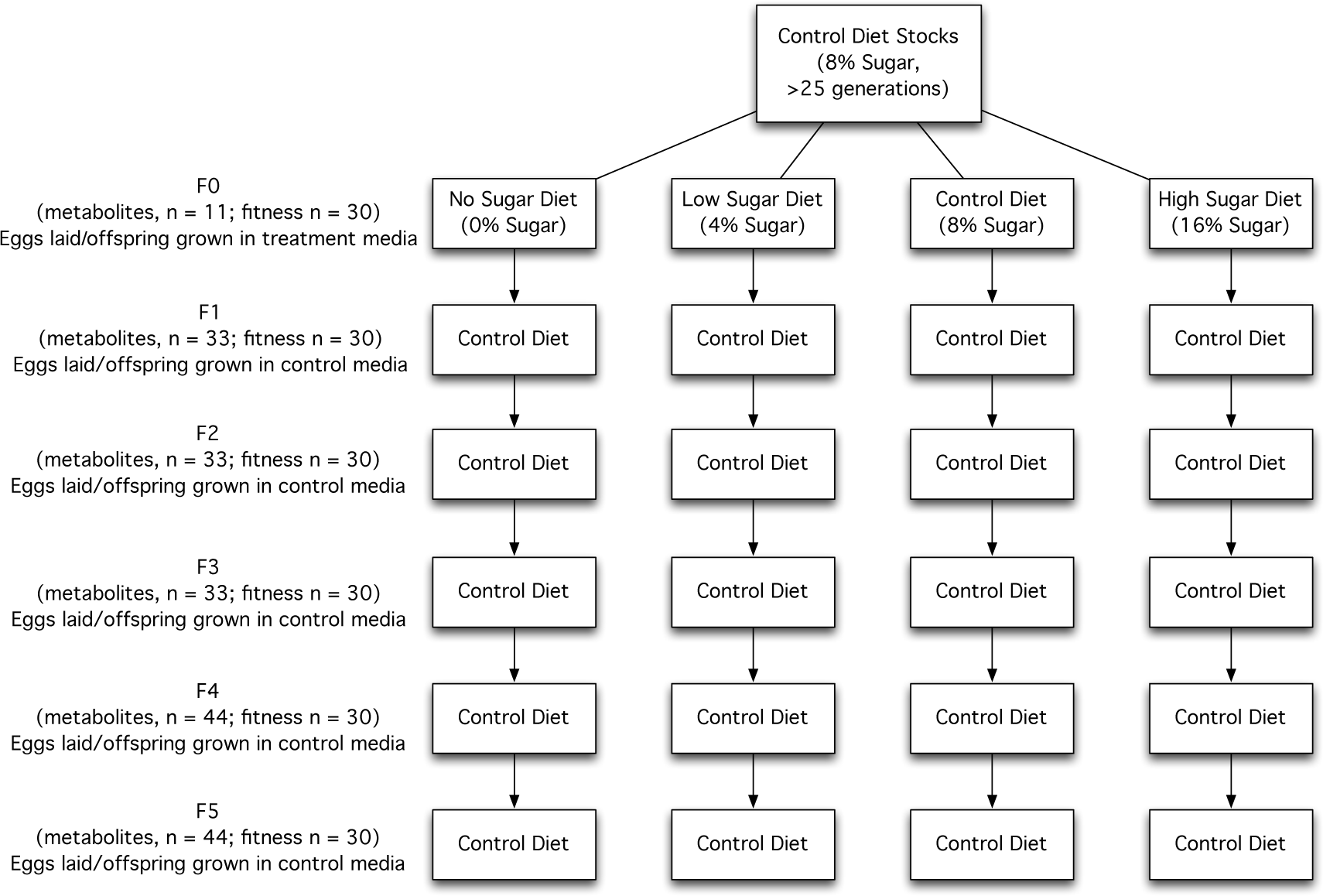
Experimental design. In the F0 generation, females and males from control stocks were exposed to one of three treatment diets (N, no-sugar diet (0% glucose); L, low-sugar diet (4% glucose); or H, high-sugar diet (16% glucose) or continued on the control diet (C, 8% sugar) for their entire lifetime. Then, all subsequent generations of offspring (F1-F5) for both treatments and controls were exposed exclusively to control diet media (C) for their entire lifetime. Whole body sugar (i.e. glucose, trehalose, and glycogen) and fat (i.e. triglycerides) concentrations, as well as lifetime reproductive fitness, were measured in all generations to assess offspring responses to ancestral dietary exposure.

**Figure 2:**
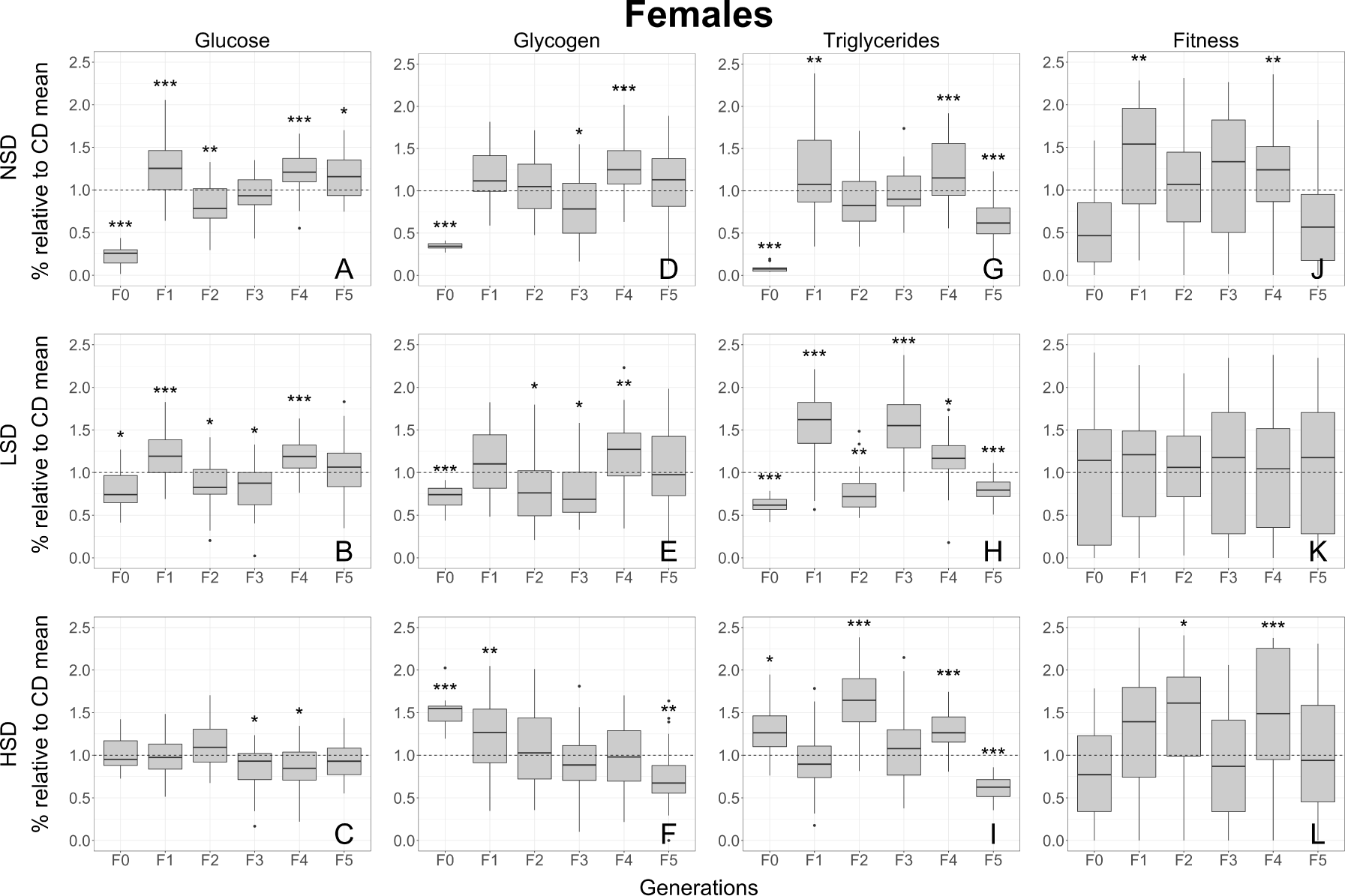
Transgenerational effects of ancestral (F0) dietary alterations on body composition and fitness in adult female flies. All data shown are relative to a control diet flies for each generation, where boxplots represent median and data quartiles for glucose, glycogen, triglyceride and fitness values relative to controls. Sample sizes for each generation are biological replicates: F0 (n=11), F1 (n=33), F2 (n=33), F3 (n=33), F4 (n=44), and F5 (n=44). Notably, given the negligible results, female trehalose values can be found in supplementary figure 1. Significance levels were corrected experiment wide using false discovery rate corrections (Benjamini and Hochberg, 1995) and are represented by *** (0.001), ** (0.01), * (0.05). Note: significant statistical differences were calculated between controls and treatments using raw values for each generation, but are displayed as ratios relative to control means for each given generation in order to better show transgenerational trends.

**Figure 3:**
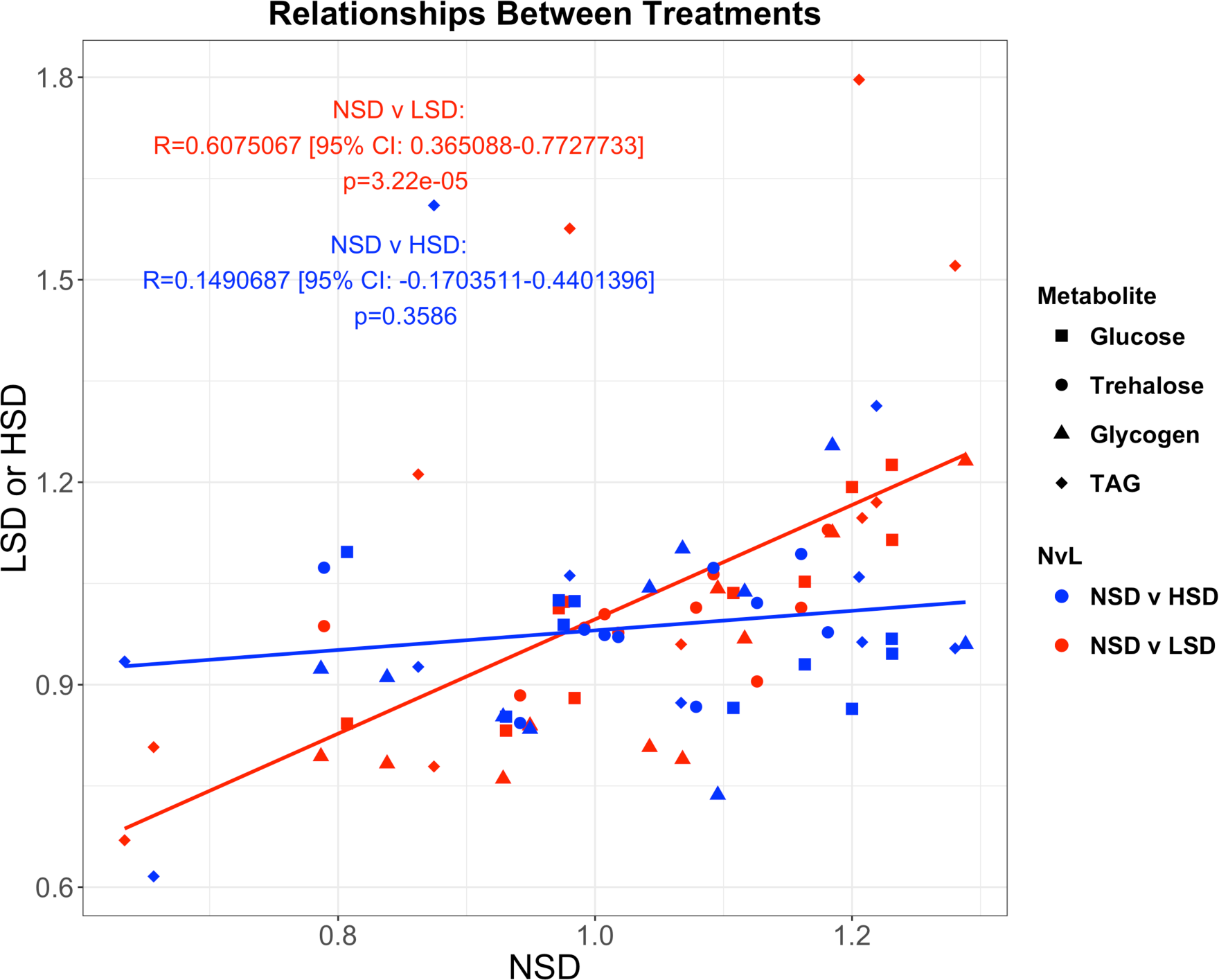
Relationships between transgenerational treatment trends. Means were calculated for each treatment and generation, from which, a Pearson’s product-moment correlation coefficient was then computed between treatments (NSD vs LSD and NSD vs. HSD) for each of the F1-F5 generations.

### Ancestral dietary treatments affect metabolite concentrations and fitness (F1-F5)

Ancestral dietary sugar exposures significantly alter metabolite concentrations for at least five generations (F1-F5) following exposure in both extreme and moderate sugar ancestral treatments (supplementary table 2). Within ancestral treatments, the direction of the metabolite concentrations vs the control mean appears to fluctuate from generation to generation, which is particularly in the case in females (figure 2). Specifically, female flies whose ancestors were exposed to the NSD and LSD treatments showed between 1-2 significant fluctuations above and below the control mean from generation to generation for glucose (figures 2A and 2B), glycogen (figure 2D and 2E), and triglycerides (figures 2G and 2H). Male flies deriving from a NSD or LSD treatment ancestors also show fluctuations around the control mean in triglyceride concentrations from generation to generation, although sugars appear to be less affected (supplementary figure 2). When significant differences are present in ancestral HSD treatment flies, the direction of metabolite concentrations vs the control mean appears to trend in a downward slope from the F0 generation to the F5 generation, with the exception of female triglycerides (figure 2I), which fluctuate similar to the NSD and LSD females.

Unlike metabolite concentration effects, fitness was completely unaffected in the LSD ancestral treatment exposure (figure 2K). However, individuals with ancestral NSD and HSD exposures had significantly more eclosed offspring in the F1 (≤0.01) and F4 (≤0.01) generations, and F2 (≤0.05) and F4 (≤0.001) generations, as compared to control flies, respectively (figures 2J and 2L).

### Relationships between transgenerational treatment trends

Generally, all metabolites in the no- and low-sugar ancestral treatment groups were positively correlated across generations (figure 3). Specifically, positive correlations were found between the no- and low-sugar ancestral treatments (r = 0.61, *p =* <0.0001). Conversely, there was no correlation between the high-sugar ancestral treatment group, and the no sugar ancestral treatment group (r = 0.149, *p =* 0.359).

### Pilot experiment

Although the pilot experiment analyzed only sugar concentrations and had smaller sample sizes, its results were significantly congruent with those of the main study (supplementary figure 3). Significant similarity between the pilot and main studies (r = 0.501, *p* = <0.0001) shows that overall patterns observed in this experiment are reproducible across experimental trials.

## Discussion

It has become increasingly clear that transgenerational phenotypic plasticity occurs (Dunn and Bale, 2011, Ayyanath et al., 2013, Herman et al., 2012, Aerts and Van Assche, 2006, Lamm and Jablonka, 2008), yet our broader understanding of these inherited and altered phenotypic traits remains largely incomplete (Moczek, 2015). In the current study, we employed a more complex experimental design approach to better understand this phenomenon. For this study, we were specifically interested in 1) the consistency of transgenerational effects from differing concentrations of the same stressor, 2) the stability of transgenerational effects over longer timescales than is frequently studied, and 3) how these responses relate to both physiology and evolution by analyzing more inclusive measurable proxies for transgenerational plasticity (i.e. body composition and fitness). Generally, we found that the ancestral treatment diets significantly influenced metabolite concentrations and fitness relative to controls for at least five generations after exposure, which influenced transgenerational phenotypes in ways that are not commonly discussed in the literature (e.g. stable, delayed and diminishing effects). Specifically, we found that males and females show different patterns of transgenerational response to the same ancestral dietary exposures, which was the case in all treatments. We also found significant relationships of response trends between no- and low-sugar ancestral treatments, yet generally no relationships between the two extreme no- and high-sugar ancestral treatments. Furthermore, we found that all ancestral treatments resulted in some degree of transgenerational fluctuations both above and below the control mean following the ancestral exposure. Interestingly, the response patterns seen from the differing treatment concentrations over longer-transgenerational timeframes have not been described previously. These findings further discussed below not only underscore the importance of transgenerational timescales and the concentration of the stressors on transgenerational phenotypic plasticity, but also give further insight into the purpose and evolutionary context of transgenerational phenotypic plasticity.

### Sex-specific differences

Our results show males and females respond significantly differently to the treatments. Sex-specific differences are not surprising in this study given that males and females have previously been shown to respond differently to environmental and heritable cues (Zambrano et al., 2005), reproductive development (Gerisch et al., 2001), dietary stress resistance (Magwere et al., 2004), and metabolism (Macotela et al., 2009). Reproductive strategies may explain the differences seen between males and females, as males use their energy stores early on for courtship and mating, and females allocate their energy and nutritional throughput for egg production throughout life, which can reach maximum output in mid-life (Cordts and Partridge, 1996). Developmental differences between males and females may also help explain the differences seen. For example, insulin-like growth factor-1 signaling (which helps regulate energy homeostasis, metabolism, and reproduction) has been shown to differ between males and females (Clancy et al., 2001, Tatar et al., 2001, Tu et al., 2002, Burks et al., 2000). In the future, transgenerational gene expression studies in males and females following ancestral treatment exposures may help better elucidate the mechanisms regulating the sex differences, and may further help us interpret transgenerational study findings.

### Relationships between treatment group trends

In a paper discussing transgenerational studies in animals, Kovalchuk (2012) questioned whether differing concentrations of a stressor can lead to similar transgenerational responses (Kovalchuk, 2012). To date, most studies documenting ancestral dietary modifications on offspring response only focus on a single extreme modification (e.g. a high-fat diet, a high-sugar diet, or a protein-restricted diet) (Zambrano et al., 2005, Benyshek et al., 2006, Dunn and Bale, 2011, Matzkin et al., 2013, Dew-Budd et al., 2016, Ost et al., 2014, Hoile et al., 2011). To better understand whether varying concentrations of the same ancestral dietary modification can lead to similar transgenerational responses, we compared the transgenerational responses of the three ancestral dietary sugar concentrations in flies.

Interestingly, although all three treatments displayed fluctuating transgenerational treatment trends, their trends were not all the same. For example, when no- and low-sugar ancestral treatment groups were compared, we found that the two treatment groups show a strong correlation between their transgenerational trends for all metabolites, suggesting similar transgenerational mechanisms. Conversely, when the opposite extreme treatment groups were compared (i.e. no-versus high-sugar ancestral treatments), there was no correlation between treatment effects (figure 3). Here, we show that differing concentrations of dietary sugar can, but do not always, produce the similar transgenerational responses.

The differences seen between the two extreme treatments may be due to the varying nature of the stress (i.e. the two treatments are regulated by, and actually lead to, completely different responses within the organism). In the case of sugar stress, starvation and low-sugar concentrations have been shown to lead to oxidative stress in *D. melanogaster*, which are regulated by the glucose homeostasis- and growth promotion-mediating *Drosophila insulin like protein* (Dilp2) (Kim and Neufeld, 2015, Post and Tatar, 2016, Rovenko et al., 2015). Conversely, high-sugar diets have been shown to lead to obesity-like phenotypes and comorbidities, which are regulated by the lipid homeostasis mediating Dilp3 in Drosophila (Kim and Neufeld, 2015, Post and Tatar, 2016, Rovenko et al., 2015). Thus, these differing responses may lead to differing transmission cues to future generations, which may help explain the patterns observed.

In a broader context, these results of differing patterns may be applicable to a number of other stressor scenarios (i.e. thermal hot versus cold stresses, pharmaceutical beneficial versus harmful dose stresses, etc.), and should be taken into consideration in future intergenerational and transgenerational studies. Thus, although the current study only begins to address Kovalchuk’s question, it highlights the importance of caution when making any broad generalizations from studies analyzing the transgenerational response to only a single modification.

### Patterns of treatment responses fluctuate relative to control from generation to generation

When analyzing offspring responses following ancestral exposure to a control and three treatment diets, we found that metabolite responses often fluctuated above and below the control mean from generation to generation. Interestingly, these fluctuations continue for several generations longer than previously believed or documented. Additionally, fitness responses were significantly altered in the two extreme (NSD and HSD) ancestral treatment groups relative to controls, where values were significantly elevated above controls in the F1 and F4, and F2 and F4, generations, respectively (Figures 2J and 2L). Given that all treatments in this study were conducted simultaneously and under identical conditions for each generation, the F0 ancestral treatment exposure differences are an obvious candidate influencing the responses seen. However, as treatment group offspring responses frequently fluctuate relative to control from generation to generation, they do not fit the interpretations commonly described in transgenerational studies.

For instance, when interpreting intergenerational and transgenerational study results, discussions frequently tend to focus on stable, delayed, or diminished inheritance. While some evidence exists for the transmission of stable effects in the literature (Remy, 2010, Rechavi et al., 2014, Ashe et al., 2012), the fluctuating responses found in our study generally do not appear to match with these findings. Conversely, delayed and diminishing responses are quite common for transgenerational studies (J Marshall and Uller, 2007, Gluckman et al., 2005, Nystrand et al., 2016, Xia and De Belle, 2016, Dew-Budd et al., 2016, Buescher et al., 2013, Dunn and Bale, 2011, Pentinat et al., 2010, Franklin et al., 2010, Walsh et al., 2015). Unfortunately, these aforementioned interpretations are frequently based on studies with short-time scales and single-exposure changes, limiting the scope of inference. For example, Benyshek and colleagues found that the direct offspring (F1) of malnourished rats showed no differences in insulin:glucose ratios from controls, yet subsequent generations (F2 and F3) had significantly increased insulin to glucose ratios, indicating a delayed transgenerational response (Benyshek et al., 2006). However, as this study, as well as many other transgenerational studies describing delayed or diminished responses, have generally only analyzed the minimum timeframes required to show transgenerational inheritance (i.e. two to three generations following exposure), they were unable to determine if the altered responses seen in the last generation(s) remained steady or continued fluctuating, and thus did not delve further into explanations or interpretations of results. Given that our study documents fluctuating responses in all three treatments for several generations beyond the minimum timeframes documented previously, and has been replicated in twice in our laboratory, our results suggest that the effects seen may be more complex than the previously discussed delayed or diminished inheritance responses.

Another less frequently used explanation for transgenerationally fluctuating responses is transgenerational compensation, where significant transgenerational effects are observed, but the signals transferred to- and the subsequent phenotype of-offspring are intergenerationally adjusted based on the interaction between the parent’s phenotype and interpreted environment (Vyssotski, 2011). For example, two recent studies found that when an individual is exposed to a stressor (e.g., protein restriction (Vyssotski, 2011, Hoile et al., 2011) and morphine treatment (Vyssotski, 2011)), that several (not one) heritable epigenetic changes were distributed over several independent loci, leading to a few similar changes, many opposite changes, and several qualitatively new changes in the offsprings’ transcriptomes from generation to generation, relative to the exposed ancestor. Although the altered gene transcription does not guarantee altered phenotype, these mechanistic findings dovetail our results. As our study documents fluctuating and opposing responses for many generations beyond F1’s direct exposure generation, the idea that parental experiences could be intergenerationally (i.e. parentally, not ancestrally) cuing offspring responses over transgenerational timeframes is possible. In this scenario, the F0 ancestral altered sugar exposure transgenerationally induces a compensatory response to try to return to homeostasis and the parent(s) determines the direction of these altered responses. However, currently there is not enough data in the literature to support this possible explanation one way or another. Thus, additional investigations could be beneficial in order to help us interpret these phenomena and understand its broader role in evolution.

## Conclusions

In conclusion, we found that ancestral diets significantly influence transgenerational metabolite concentrations and fitness for at least five generations following exposure, despite the conformity to a control diet in the F1-F5 treatments. Given that fly responses were simultaneously analyzed for both sexes in all three treatments for each generation, we were able to show that transgenerational responses differed between sexes, and that transgenerational response trends can be similar or different to each other depending on the treatment. Additionally, as responses were measured for several generations longer than seen in most transgenerational studies previously, we were able to observe that fly responses to the ancestral treatment diet can fluctuate above and below control means for at least five generations following the inducing exposure. Notably, the response fluctuations observed relative to controls from generation to generation shed doubt on previous study interpretations, including the common assumption of universal transgenerational response stability. Finally, as we conducted both a pilot and main study with the same study design, we have been able to replicate our findings of fluctuating transgenerational responses. Overall, the patterns found in the current study do not fit with any of the commonly proposed explanations, although a newer, less common explanation of transgenerational compensation may show promise in the future. Additionally, though not discussed here, developmental processes and additional external factors (i.e. host-microbe interactions) may be playing a role in the patterns observed (Laland et al., 2015). Thus, studies analyzing mechanisms and external factors influencing transmission of the patterns seen here deserve further attention. Taken together, this study not only highlights the need for additional studies analyzing multiple ancestral dietary modifications and longer transgenerational timeframes, but also additional phenotypic and mechanistic data to help explain the patterns observed. Furthermore, by increasing the number of more complex studies, we may begin to better understand the purpose and evolutionary value of transgenerational plasticity and inheritance.

## Materials and Methods

### Fly stocks

A single phenotypically inbred wild type population of (Canton-S-brn) *D. melanogaster* was obtained from Drosophila Genetic Research Center (Kyoto DGRC #109019), Japan. Stock flies were raised and maintained in glass vials in a standard yeast/glucose diet (4% yeast, 8% dextrose, 1% agar, 0.4% propionic acid, 0.3% butyl-*p*-hydroxy benzonate) at 25^º^C and 60% relative humidity under 13h:11h light:dark cycles. Prior to this study, flies were maintained with a control diet for >25 generations.

### Exposure diets and experimental design

In the first generation (F0) of this study, wild-type stock flies were exposed to one of four diets: no-sugar diet (0% sugar, NSD); low-sugar diet (4% sugar, LSD); control diet (8% sugar, CD); and high-sugar diet (16% sugar, HSD); where all other media ingredients beyond sugar stayed constant (1% agar, 4% yeast, 0.7% preservative, RO water). Flies that were sacrificed for metabolic analyses were exposed to their respective treatment media from oviposition to death. Flies used to create the next generation were exposed from oviposition until six days old, at which time they were transmitted to new vials containing control diet media for creation of the F1 generation. P1 (F0) files remained in the control diet media in order to lay eggs for three days, at which time they were removed and euthanized. For all subsequent generations (F1-F5), each treatment group was exposed exclusively to control diet media from oviposition to death and all treatments were done simultaneously for each generation (figure 1); thus, any resulting phenotypic between-group differences for a given generation resulted from ancestral exposure differences. Notably, the density of flies grown in each vial from stocks and treatments for all generations were controlled by mating six females with four males for 72 hours, which was determined as the optimum mating strategy for our targeted population size prior to experimentation.

### Sample collection for metabolic analysis

For all generations, virgin flies, which had been exposed to their respective generation’s treatment for their entire lives, were collected within eight hours of eclosion and stored in sex-separated vials containing fresh media, which corresponded to the media they were reared in. Notably, to prevent psuedoreplication, each sample was maintained in its own vial separate from other samples throughout their life. At seven days old, these offspring were starved for 24 hours in order to clear guts of biasing media contents. After 24 hours of starvation, pooled samples of four flies were weighed to the nearest 0.1 mg and processed for metabolite measurements. For metabolites, sample sizes for each generation were as follows: F0 (n=11 pooled samples of 4 flies per sample), F1 (n=33 pooled samples of 4 flies per sample), F2 (n=33 pooled samples of 4 flies per sample), F3 (n=33 pooled samples of 4 flies per sample), F4 (n=44 pooled samples of 4 flies per sample), and F5 (n=44 pooled samples of 4 flies per sample). Notably, sample sizes for each generation were determined using a power analysis based on our pilot study.

### Sugar quantification

Pooled whole fly samples were homogenized in ice-cold acetate buffer (pH 5.6), incubated at 95^º^ C for 20 minutes to prevent degradation, and centrifuged at 12,000 rpm for 2 minutes. The resulting supernatant was collected for glucose, trehalose, and glycogen analysis. Trehalose and glycogen samples were treated with trehalase (0.25 units/mL) and amyloglucosidase (5 units/mL), and incubated for 12 hours at 37^º^C and 60^º^C, respectively. Resulting glucose levels for three sugars were analyzed using Glucose Assay Reagent (Sigma GAHK20), where samples and standards were randomized on the plate(s). For each generation, standards were freshly made via serial dilution of a concentrated stock for each sugar. Trehalose and glycogen concentrations were then calculated by first doubling absorbance values in order to correct for enzyme dilution and then subtracting out the amount of free glucose from each sample. Each sugar’s absorbance was then compared to a sugar-specific standard curve. Notably, samples were normalized to weight (Toth et al., 1993, Hebert and Keenleyside, 1995).

### Lipid quantification

#### Extraction

Pooled samples were homogenized in 200 µL ice-cold methanol containing internal standards using a Physcotron Handy Micro Homogenizer. Internal standards contained triheptadecanoin, a heavy triglyceride compound not found in nature (Larodin Fine Chemcials). Following homogenization, 400 µL methyl-tert-butyl ether (MTBE) was added to each sample and samples were shaken for seven minutes at 1100 rpm. Next, 100 µL HPLC-grade H_2_O was added and samples were shaken at 4^º^ C for 30 seconds at 1000 rpm. Samples were then centrifuged at 2000 rpm for five minutes. Finally, 200 µL of the top layer (MTBE containing lipids) was transferred to a new glass insert, speed vacuumed to dryness, and stored at -20^º^ C until analysis.

#### Analysis and quantification of Lipids using UHPLC-MS

For analysis, dried samples were resuspended in 150 µl of toluene and sonicated for 10 minutes. Then, 10 µL of resuspended sample was added into 90 µL methanol, creating a ten fold dilution, which was sonicated for 10 min. This resuspension procedure was automated using a PAL Combi-xt autosampler. The autosampler syringe was washed with 400 µl toluene and 200 µl methanol between samples. For each sample, three µl of the 10-fold dilution was injected into a Waters ACQUITY UPLC Class-I in tandem with a Waters SYNAPT G2-S High Definition Mass Spectrometer equipped with Ion Mobility. Lipids were separated in an ACQUITY UPLC CSH C18 1.7 µm 2.1 x 100 mm analytical column at 400 µL/min, 60^º^ C. A separation gradient was used to separate compounds and comprised of two solvents (A and B). Solvent A was comprised of a 60:40 Acetonitrile:Distilled water (10 mM Ammonium Formate + 0.1% Formic Acid) solution, and Solvent B was comprised of a 90:10 2-Isopropanol:Acetonitrile (10 mM Ammonium Formate + 0.1% Formic Acid) solution. The gradient shift began with 85% solvent A and 15% solvent B, shifting to 40% solvent A and 60% solvent B in three minutes, then to 28% solvent A and 72% solvent B in 0.5 minute, then to 20% solvent A and 80% solvent B in 4.5 minutes, then to 0% solvent A and 100% solvent B in one minute, and held at 99% solvent B for two minutes. The column was then equilibrated for one minute at 15% solvent B, followed by a post-separation washing gradient of 99% solvent B for two minutes, and a final equilibration at 15% solvent B for 2 minutes. Total run time was 17 minutes. Autosampler solvents were comprised of 60:40 Acetonitrile:Distilled water, which was used for aspirating and loading sample into sample loop, and 90:10 2-Isopropanol:Acetonitrile (0.1% Formic Acid) for washing the needle to avoid carryover between samples. Mass spectrometer used a LockMass solution of Leucine/Enkephalin 2 pmol/ml in 50% Acetonitrile (0.1% Formic Acid) infused every 30 seconds for automatic mass correction during acquisition time. Mass spectrometer settings were as follow, 2.0 kV spray voltage, cone voltage 30 V, desolvation temperature 400^º^C, desolvation gas 900 L/Hr, source temperature 120 ^º^C, acquisition range from 50 to 1700 m/z, scan rate 10 hz, acquisition mode MSe (Independent Data Acquisition), high resolution 35,000 FWHM, continuum mode, quad profile automatic, collision energy was 6 V for low energy (Collision Trap), and ramped from 20 to 40 V in high energy mode. Mass spectrometer was calibrated with Sodium Formate 500 mM in water.

Acquisition of mass spectrometric data was done using Waters MassLynx v4.1. Chromatographic data was processed using MzMine2 open-source software, for mass correction (using acquired lock mass data), alignment, normalization, deconvolution of high energy data (MSe), isotope grouping, peak picking and peak identification based on high energy fragmentation using Lipid Maps database (18 Mar 2014 version). Following peak identification, possible metabolic species were listed and individual compounds were manually assigned from this list based on isotope similarity, compound score (as provided by software), and expected retention times. The total sum of all identified triglycerides were then divided into an internal standard, which was added to the sample prior to processing and provided relative lipid concentrations for each sample.

### Lifetime reproductive fitness

Fitness represented the total nuFmber of successfully eclosed offspring produced by a single female from eclosion until death (n = 30 for each treatment and generation). Briefly, upon eclosion, one female and one non-sibling male were placed into a vial containing one of the dietary treatments described previously (i.e. parents of F0 (P0) were mated in a NSD, LSD, CD, or HSD; P1 = F0 and were mated in CD). To make sure that female reproduction was not limited by male quality, a new male was transferred into each vial every second week, or immediately if escaped during handling or found dead. Flies used for this fitness study were moved to new vials twice per week in order to prevent overcrowding and to reduce counting errors. Twice per week, the number of eclosing flies were counted from each vial and tallied over the course of the female’s lifetime.

### Statistical analyses

Data were analyzed using R statistical software (version 3.2.1). In order to help account for the complexity of multiple factors influencing responses, linear regressions were used to calculated residuals for the multivariate model, where fixed variables comprised of ancestral dietary treatment, sex, and total pooled fly weight for each endpoint (metabolite and fitness responses) within a given generation. Notably, in order to better show transgenerational trends, we compared raw treatment values to control values within generations and display our data as relative differences between controls and treatments for each generation. Prior to analysis, linear model assumptions were validated. In order to account for Type I errors associated with multiple comparisons, False Discovery Rate (FDR) corrections were conducted using the Benjamini-Hochberg procedures (Benjamini and Hochberg, 1995) to control experiment-wise error rates.

To test for relationships between transgenerational trends of the three treatments, means were calculated for each treatment and generation. From this, a Pearson’s product-moment correlation coefficient was then computed between treatments for each of the F1-F5 generations. Detailed sttatistical analysis and results are included in the supplementary materials. All data, statistics, and tables can be found at: https://github.com/cemborski/DynamicTransgenerationalPlasticity_SupplementaryMaterials.git (Emborski, 2017)

### Pilot Study

Prior to this main study, a pilot experiment was conducted. As the pilot was the basis for the main study, sugar metabolites were analyzed using the same methodology described herein for the same three treatments and control over six generations (*i.e.* F0-F5), but did not include fat metabolite analysis. Analysis of differences between treatments and sexes were conducted for each generation using the same methodology as the main study outlined previously, where the data, statistics, and results can be found in the supplementary documentation. For each generation, treatment, and sex, eight samples were collected and analyzed (vs. n=11 (F0), n=33 (F1-F3), and n=44 (F4-F5) in the main study). To test for a relationship between the main study and pilot study transgenerational trends, means were calculated for each treatment and generation. From this, a Pearson’s product-moment correlation coefficient was then computed between the two studies for each of the F0-F5 generations.

## Acknowledgements

We are deeply grateful to the OIST Mass Spec Center, and particularly Alejandro Villar Birones for acquiring the LC/MS data and extensive technical consultation. Additionally, we would like to thank Mandy Tin, Maggi Mars Brisbin, Ken Meacham, Stephen Emborski, Marco Tsui, and Takakazu Yokokura for their assistance, guidance, and support throughout this project.

## Competing Interests

We have no competing interests.

## Funding

Funding for this study has been provided by the OIST subsidy budget.

## Availability of data and materials

All data, statistics, and tables analyzed for this publication can accessed via our github repository at: https://github.com/cemborski/DynamicTransgenerationalPlasticity_SupplementaryMaterials.git

## Authors Contributions

Carmen Emborski conceived of the study, carried out the lab work, conducted the data analysis, and drafted the manuscript. Alexander Mikheyev participated in the design of the study, the statistical analysis, the writing of the manuscript, and obtained funding. All authors gave final approval for the publication.

